# Glucocorticoid–endocannabinoid crosstalk in the ventrolateral periaqueductal gray (vlPAG) promotes pain resolution

**DOI:** 10.64898/2026.04.25.720827

**Authors:** B. Coutens, C.A. Bouchet, I. Gvon, L.C. Patti, C.M. De Anda Gamboa, D.C. Jewett, Je. Klawitter, J. Klawitter, M.M. Heinricher, S.L. Ingram

## Abstract

Inflammation is a primary response to injury. Here we show that inflammation plays a critical role in engaging the endocannabinoid system in the ventrolateral periaqueductal gray (vlPAG) to activate the descending pain modulatory circuit to inhibit pain. Inflammation-induced increases in corticosterone activate glucocorticoid receptors to increase synthesis of 2-arachidonylglycerol (2-AG). Retrograde transmission of 2-AG stimulates presynaptic cannabinoid 1 receptors to inhibit GABA release in the vlPAG, producing anti-hyperalgesia. Conversely, blocking both glucocorticoid and cannabinoid receptor activity impairs recovery from hyperalgesia, highlighting the beneficial role of endocannabinoid signaling in pain resolution. However, this system is tightly regulated and over-stimulation of glucocorticoid receptors with corticosterone results in cannabinoid 1 receptor desensitization. In addition, cannabinoid receptors are more susceptible to desensitization in inflamed rats and rapidly desensitize in response to exogenous cannabinoid receptor agonists. Thus, there is a narrow therapeutic window for cannabinoid drugs in the context of inflammatory pain. These findings indicate that cannabinoid agonists should be used with caution in the context of inflammation to avoid CB1R desensitization, and that exploiting glucocorticoid-endocannabinoid interactions is a promising strategy to optimize cannabinoid-based therapies for inflammatory pain.

## Introduction

Cannabis has been used to manage pain for millennia, but the current push to legalize marijuana and other cannabinoids has increased use substantially in the past 5 years [1]. Randomized, controlled clinical trials of cannabinoids for treatment of chronic pain show a statistically significant, albeit small, change in patient reports of pain [2-5], although not all studies find clinical benefit [6]. In preclinical studies, cannabinoids are reported to reduce acute pain responses, as well as reversing hyperalgesia associated with chronic pain [7-10], with greater efficacy at producing anti-hyperalgesia [11]. Thus, there is a mismatch between preclinical and clinical efficacy that highlights a gap in understanding cannabinoid mechanisms [12]. There is nonetheless agreement that effects of exogenous cannabinoids on pain are mediated through activation of cannabinoid 1 receptors (CB1Rs) [7].

By contrast with the intensive study of exogenous cannabinoids, the role of *endogenous* cannabinoids in modulating pain has been much less well studied. From a drug-development perspective, the primary approach has been to raise endocannabinoid levels using blockers or knockout of degradative enzymes, primarily monoacylglycerol lipase (MAGL) and fatty acid amide hydrolase (FAAH) [13-18]. These studies indicate that blocking MAGL or FAAH can be effective at inhibiting acute and chronic pain in pre-clinical studies [13-18], yet clinical trials have not proven this approach to be effective in patients with chronic pain [19].

A different approach, administration of a CB1R antagonist, can be used to determine whether pain is subject to ongoing CB1R activation by endocannabinoids, and a small number of studies using the CB1R antagonist rimonabant (RIM) have provided evidence in support of endocannabinoid tone under some conditions [20-22]. A better understanding of endocannabinoid regulation will therefore be needed to harness the potential of targeting this system for development of novel pain therapeutics. Here we consider the role of endocannabinoids in limiting hypersensitivity and promoting recovery in animals subjected to persistent inflammation.

One possible mechanism for endocannabinoid recruitment in persistent pain states is via elevated cortisol and other glucocorticoids. These are elevated in inflammatory conditions [23,24]. Cortisol (in humans) and corticosterone (CORT, in rats) act at the glucocorticoid receptor (GR) to stimulate synthesis of the endocannabinoid 2-AG leading to activation of CB1Rs [25-28]. GRs are widely expressed in pain-related brain regions [29], including the ventrolateral periaqueductal gray (vlPAG), a brain area known to play a rôle in the antinociceptive actions of opioids and exogenous cannabinoids [30]. Direct application of exogenous cannabinoids in the vlPAG leads to analgesia [31,32]. In addition, endocannabinoids are synthesized and released in this region [13,33-36], and have documented effects on vlPAG circuits [13,37,38]. Moreover, inflammation and increased CORT levels promote endocannabinoid release and activation of CB1Rs in the vlPAG [27,34]. Viewed as a whole, these lines of evidence raise the possibility that glucocorticoids and endocannabinoids work together in this region to modulate pain.

Here we hypothesized that endocannabinoids contribute to resolution of hypersensitivity in persistent inflammation. We find this occurs via elevated CORT, which activates GRs and drives the synthesis of endocannabinoids within the vlPAG. The resulting CB1R-mediated engagement of vlPAG circuits in turn contributes to resolution of inflammation-induced behavioral hypersensitivity. GR and CB1R antagonists were used to assess endocannabinoid and glucocorticoid tone in the vlPAG and their role in behavioral measures of pain. Whole-cell patch-clamp electrophysiology was employed to directly measure endocannabinoid activation of CB1Rs in the vlPAG. Our findings delineate a finely tuned regulatory interplay between glucocorticoid and endocannabinoid signaling within the vlPAG, thereby providing a mechanistic framework to inform and enhance endocannabinoid-based strategies for pain management.

## Materials and Methods

*Detailed materials and methods can be found in Supplementary Materials*.

### Animals

Adult male and female Sprague Dawley rats (3-11 weeks old) were used for all experiments. All procedures were performed in accordance with the Guide for the Care and Use of Laboratory Animals as adopted by the Institutional Animal Care and Use Committee of the University of Colorado Anschutz Medical Campus.

### Inflammation

Complete Freund’s Adjuvant (CFA; 1 mg/ml, 0.1 ml; Sigma-Aldrich) was injected subcutaneously into the plantar surface of the right hindpaw. [39]

### Drugs

WIN55,212-2 (WIN), SR141716A (RIM), Corticosterone (CORT), 11b-(4-dimethyl-amino)-phenyl-17bhydroxyl-17-(1-propynyl)-estra-4,9-dien-3-one (RU486) were dissolved in dimethyl sulfoxide (DMSO), aliquoted, and stored at -20°C. NBQX was dissolved in milliQ water, and stored at 4°C. Compound101 (Cpd101) was first dissolved in a small amount of DMSO (10% of final volume), sonicated, then brought to its final volume with 20% 2-hydroxypropyl)-b-cyclodextrin (HPCD) and sonicated again to create a 10 mM solution. PKA inhibitor (PKI) was used directly in the internal solution in recording electrodes at 0.2 µM.

### vlPAG slice preparation

Slices containing the vlPAG were prepared as previously described [34,40]. Rats were deeply anesthetized with isoflurane (McKesson, Irving TX, USA), and the brain was rapidly removed and placed in ice-cold aCSF cutting buffer containing the following (in mM): 126 NaCl, 21.4 NaHCO3, 22 dextrose, 2.5 KCl, 2.4 CaCl2, 1.2 MgCl2, and 1.2 NaH2PO4 (300 mOsm). Slices containing the vlPAG were cut to a thickness of 220 µm on a vibratome (Leica Microsystems, Deerfield IL, USA) and were transferred to a holding chamber maintained at 32°C. Slices were oxygenated with 95% O2/5% CO_2_ until transfer to the recording chamber and superfused with oxygenated aCSF maintained at 32°C.

### Whole-cell patch-clamp recordings

Voltage-clamp recordings (holding potential, -70 mV) were made in whole-cell configuration using an amplifier (MultiClamp 700B, Molecular Devices), sampled at 2 kHz, and digitized at 5 kHz with the Axon Digidata 1550B (Molecular Devices, USA) using Clampex 11.0.3 software (Molecular Devices, USA). Patch-clamp electrodes were pulled from borosilicate glass (diameter, 1.5 mm; WPI). Pipettes (2.5 - 4 MΩ) were filled with an intracellular solution containing (in mM): 140 CsCl, 10 HEPES, 4 MgATP, 3 NaGTP, 1 EGTA, 1 MgCl2, and 0.3 CaCl2 (pH 7.3, 290–300 mOsm). QX314 (100 µM) was added to the internal solution for evoked IPSC (eIPSC) experiments to reduce action potentials in the recording cell. Access resistance was continuously monitored. Recordings in which access resistance changed by 20% during the experiment were excluded from data analysis.

### Depolarization-induced suppression of inhibition

After obtaining stable eIPSCs, a protocol for DSI collected 2-3 eIPSCs for baseline measurement, followed by a brief depolarizing step (5 s at +20 mV; [41]) before returning to the holding potential. eIPSCs were evoked at 0.2 Hz for 60 s following the depolarizing step, and normalized to the average baseline eIPSC amplitude. Cells were grouped into “DSI” or “No DSI” with DSI defined as a minimum of 10% inhibition of eIPSCs.

### Endocannabinoid analysis

Quantification of 2-AG and AEA level were performed in the vlPAG, the RVM and the injured paw tissue of naïve and CFA-treated rats. Tissues were removed, quickly frozen in liquid nitrogen and stored at -80ºC degrees until analysis as previously described [42-44].

### Measurement of plasma CORT levels

Plasma CORT levels were determined using trunk blood immediately after anesthesia at the time of slicing for electrophysiological recordings in the morning to avoid variation due to circadian rhythm. All samples were analyzed using the commercially available enzyme-linked immunosorbent assay.

### Behavioral studies

Mechanical nociception was assessed using an electronic von Frey (Ugo Basile, Gemonio, Italy). Responses from each hind paw were measured three times each, and the mean for each paw was calculated as the paw withdrawal threshold (PWT). All measurements were performed by the same person, blinded to treatment. Thermal nociception was assessed using the Hargreaves apparatus (Plantar Test, Ugo Basile, Gemonio, Italy), which measures paw withdrawal latency (PWL) to radiant heat stimuli. The radiant heat source was applied to the center of the plantar surface of each hind paw with 2 min intervals between each application, and PWL determined as time to retraction or licking of the hind paw with a cut-off of 25 seconds. All trials were performed three times for each hind paw, and the average for each hind paw was calculated as the PWL. All measurements were performed by the same person who was blinded to the treatment.

### vlPAG microinjections

Unilateral microinjection of RU486 (3 µg/kg dissolved in 10% DMSO in saline, 0.4 µl in one minute) into the vlPAG was performed using a stereotaxic apparatus (NeuroSTAR, Germany). Animals were subjected to the behavioral testing protocols 15, 30 and 60 min post injection.

### Statistical analysis

All analyses were conducted in Graphpad Prism (Graphpad Software). Values are presented as the mean ± SEM, and all data points are shown in bar graphs to illustrate variability. Statistical comparisons were made using t-test or ANOVA and post-hoc analyses, as appropriate. In all summary bar graphs for electrophysiology experiments, each dot represents an individual cell while the numbers in the bars represent the animal number. Statistical significance was defined as p < 0.05.

## Results

### CFA inflammation and hyperalgesia are associated with increased endocannabinoid levels in the PAG and the site of inflamation

Local injection of CFA in the hindpaw is an established model of persistent inflammation and behavioral hypersensitivity [45-47]. CFA treatment resulted in increased sensitivity to mechanical and thermal stimulation of the inflamed hindpaw that lasted at least 7 days, with recovery to baseline by 21 days post-injection (Fig. 1A).

**Figure 1.**
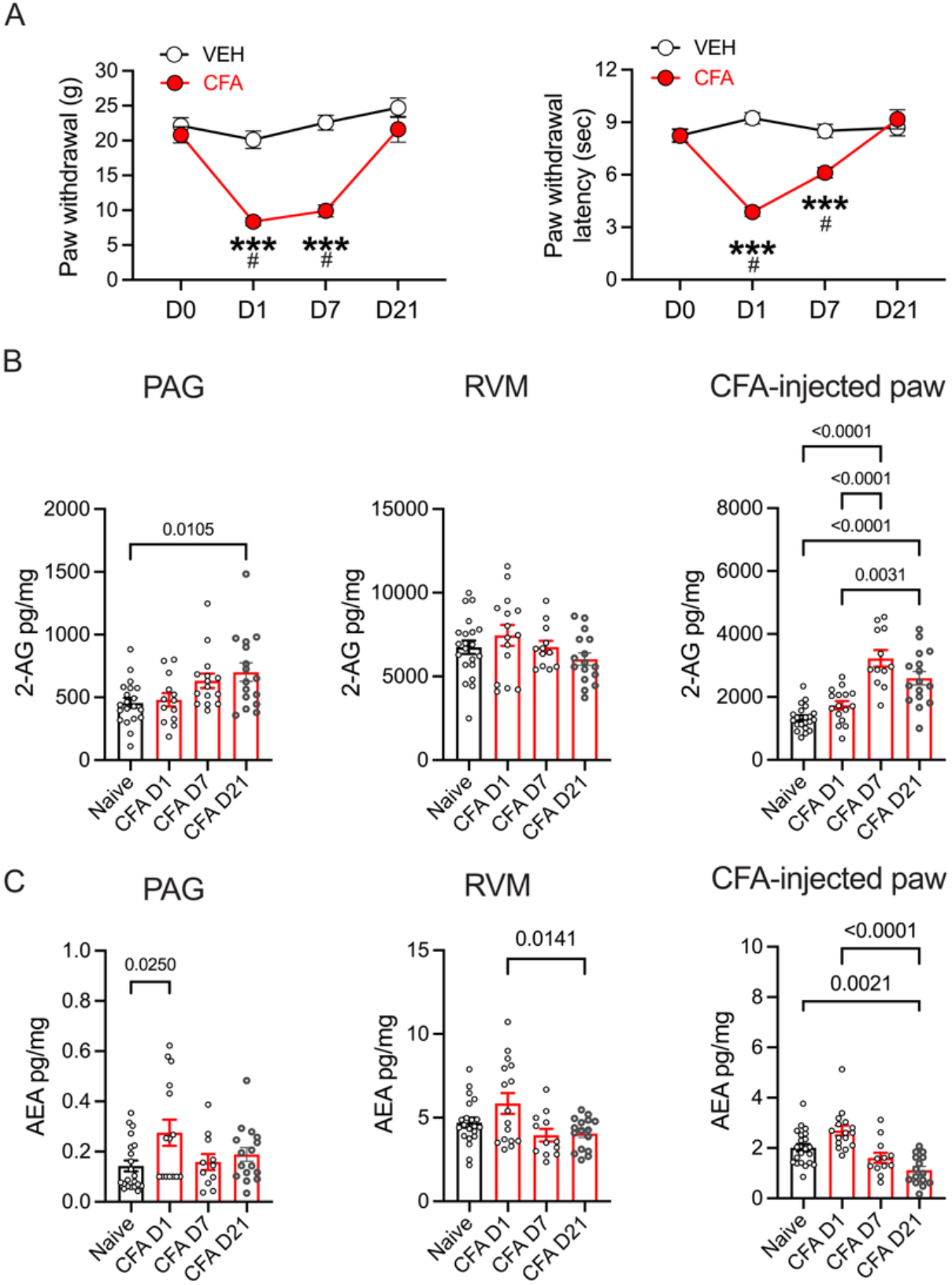
CFA inflammation induces hyperalgesia and changes in endocannabinoid levels. A. Paw withdrawal measurements using von Frey fibers show the time course of CFA-induced hyperalgesia across time (2-way RM ANOVA, Time X Treatment; F_3,66_ = 16.96, p < 0.0001, Tukey’s multiple comparisons, **** CFA compared to VEH, p < 0.0001; # Compared to D0, p < 0.05, N = 12 (6 male/6 female rats per treatment). Paw withdrawal latencies to thermal stimulation using the Hargreaves test across time (2-way RM ANOVA, Time X Treatment; F_3,66_ = 23.66, p < 0.0001, Tukey’s multiple comparisons, ****CFA compared to VEH, p < 0.0001; # Compared to D0, p < 0.05, N = 12 (6 male/6 female rats per treatment). A 3-way ANOVA showed no significant behavioral differences between males and females on the von Frey test (Sex X Treatment: F_(1, 20)_ =0.24, p = 0.63) or Hargreaves test (F_(1, 20)_ = 0.87, P=0.36). B. Levels of 2-AG measured in brain tissue (PAG: One way ANOVA, F_3,59_ = 3.35, p = 0.02, RVM: One way ANOVA, F_3,60_ = 1.55, p = 0.21) and CFA-injected paw (One way ANOVA, F_3,61_ = 24.96, p < 0.0001). Tukey’s multiple comparisons on graphs. C. Levels of AEA measured in brain tissue (PAG: One way ANOVA, F_3,59_ = 3.08, p = 0.03, RVM: One way ANOVA, F_3,61_ = 4.35, p = 0.0077) and CFA-injected paw (One way ANOVA, F_3,60_ = 13.90, p < 0.0001). Tukey’s multiple comparisons shown on graphs.

We examined levels of two primary endocannabinoids, 2-AG and AEA, in the PAG and its primary output target, the RVM, and in the inflamed hindpaw. In parallel with behavioral hypersensitivity, there was a sustained (> 21 days) increase in 2-AG levels in both the PAG and the inflamed hindpaw, but not in the RVM (Fig. 1B). By contrast with 2-AG, AEA levels were elevated in the PAG only on the first day post-CFA, and in fact decreased in the RVM and paw over time (Fig. 1C).

### CB1R antagonist reveals endocannabinoid regulation of behavioral hypersensitivity

Cannabinoids reduce pain and hypersensitivity via CB1Rs [17,47,48]. The observation that 2-AG was elevated following CFA treatment and throughout the time course of hypersensitivity raises the possibility that this endocannabinoid serves to limit hypersensitivity in persistent inflammation and/or promote recovery. We tested the possibility that hypersensitivity would be enhanced by CB1R blockade using systemic administration of the CB1R antagonist rimonabant (RIM, 3 mg/kg IP, once daily for up to 10 days). Hypersensitivity was potentiated at 7 and 10 days post-CFA, with lower thresholds for mechanical stimuli and shorter response latencies for thermal stimuli in rats given daily RIM compared to vehicle injections (Fig. 2B). Moreover, in separate groups of animals, RIM given only once, at a time point at which hypersensitivity had returned to baseline (21 days for mechanical stimulation, 10 days for thermal stimulation), reinstated hypersensitivity (Fig. 2C,D). Neither RIM nor vehicle altered thresholds in naïve animals (Suppl. Fig. 1A,B). Notably, RIM had no effect on day 1, the time of maximal hypersensitivity at which AEA, but not 2-AG, was elevated in vlPAG. These findings indicate that ongoing activation of CB1R, likely reflecting increased levels of 2-AG, limits behavioral hypersensitivity and contributes to recovery.

**Figure 2.**
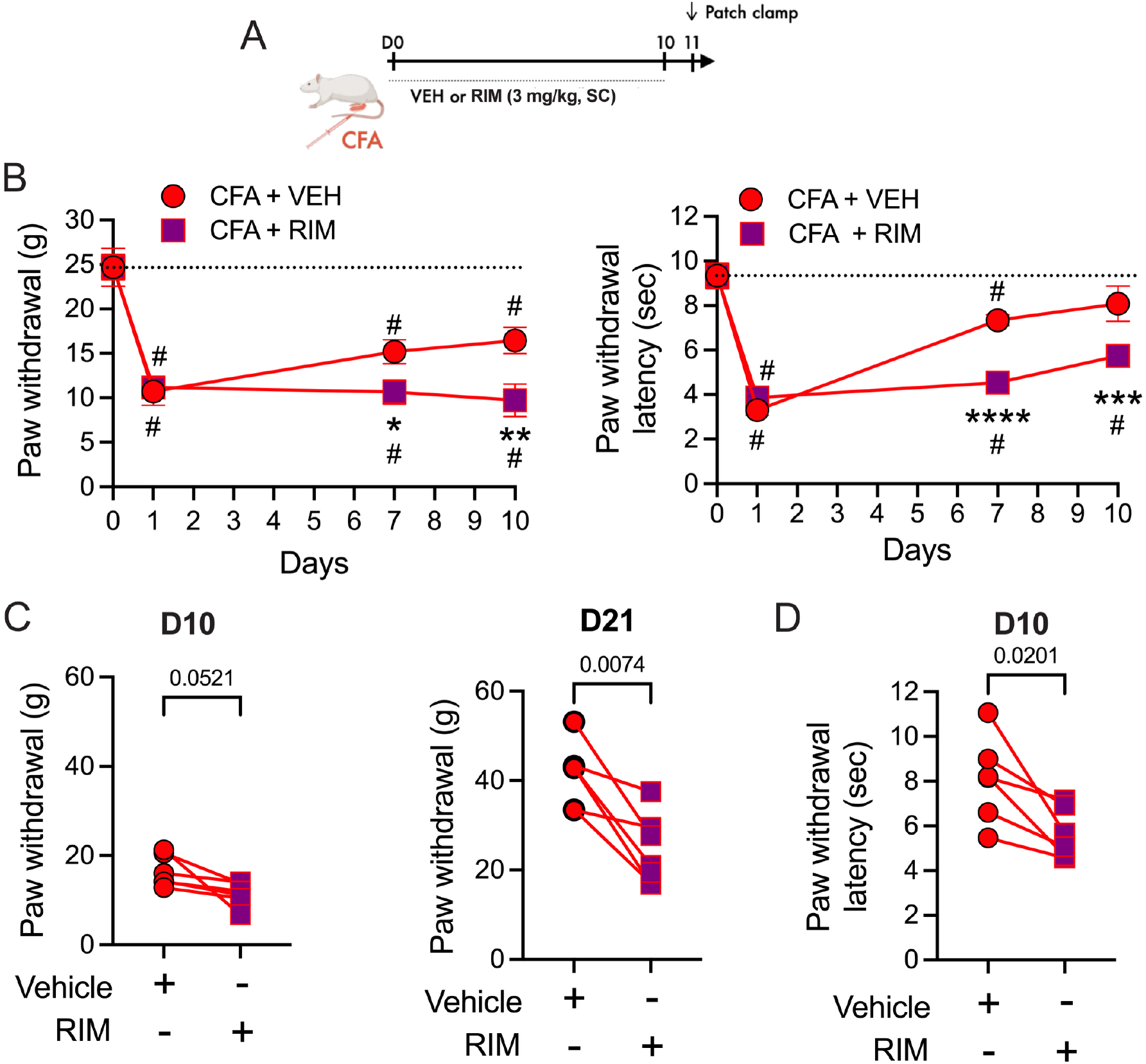
CB1R antagonist reverses eCB anti-hyperalgesia. A. Schematic showing daily injections (SC) of vehicle (VEH) or rimonabant (RIM: 3mg/kg) for 10 days after CFA injections into a hind-paw at D0. B. Behavioral measurements were made on Day 0 prior to CFA injection, and on days 7 and 10 one hour after injections. *Von Frey:* Two way RM ANOVA: Time X Treatment, F_3,30_ = 3.70, p = 0.022, Tukey’s multiple comparisons on graph, CFA compared to VEH, *p = 0.048, **p = 0.0043; # Compared to D0, p < 0.05. *Hargreaves*: Two way RM ANOVA: Interaction, F_3,30_ = 6.05, p = 0.0024, Tukey’s multiple comparisons on graph, CFA compared to VEH, ***p = 0.0007, ****p < 0.0001; # Compared to D0, p < 0.05. C. Rats treated with CFA/VEH injections received a single injection of RIM (3 mg/kg, SC) on Day 10 and behaviors were tested 30 minutes after RIM injection (Von Frey: paired t-test: t_5_ = 2.54, p = 0.0521). A different group of animals were treated with CFA for 21 days and tested with VEH followed by RIM (3 mg/kg, SC; Von Frey: One way RM ANOVA, F_2,11_ =8.164, p = 0.0067, Tukey’s multiple comparisons on graph; only post-VEH and post-CFA are shown on graph as VEH had no effect from baseline, p = 0.61). D. Thermal thresholds were significantly decreased by RIM on D10 (Hargreaves: paired t-test: t_5_ = 3.36, p = 0.020). The within-rat response is denoted by a line.

### Endocannabinoid function in the vlPAG is enhanced, and regulated, by CORT in persistent inflammation

Given the evidence that endocannabinoids limit behavioral hypersensitivity and promote recovery, we next probed endocannabinoid function in the vlPAG during inflammation using the DSI protocol [49-51] in whole-cell patch clamp recordings from vlPAG neurons in slice. We focused on day 7 and day 21, since the behavioral studies using the CB1R antagonist indicated that endocannabinoid function was behaviorally relevant at these time-points. Compared to slices from naïve rats, the DSI response was prolonged in slices from CFA-treated animals. DSI lasted approximately 30 s in naïve controls, but lasted at least 60 s in CFA-treated animals (Fig. 3A,B). This finding reinforces the conclusion from the behavioral and biochemical studies that endocannabinoids are released in persistent inflammation.

**Figure 3.**
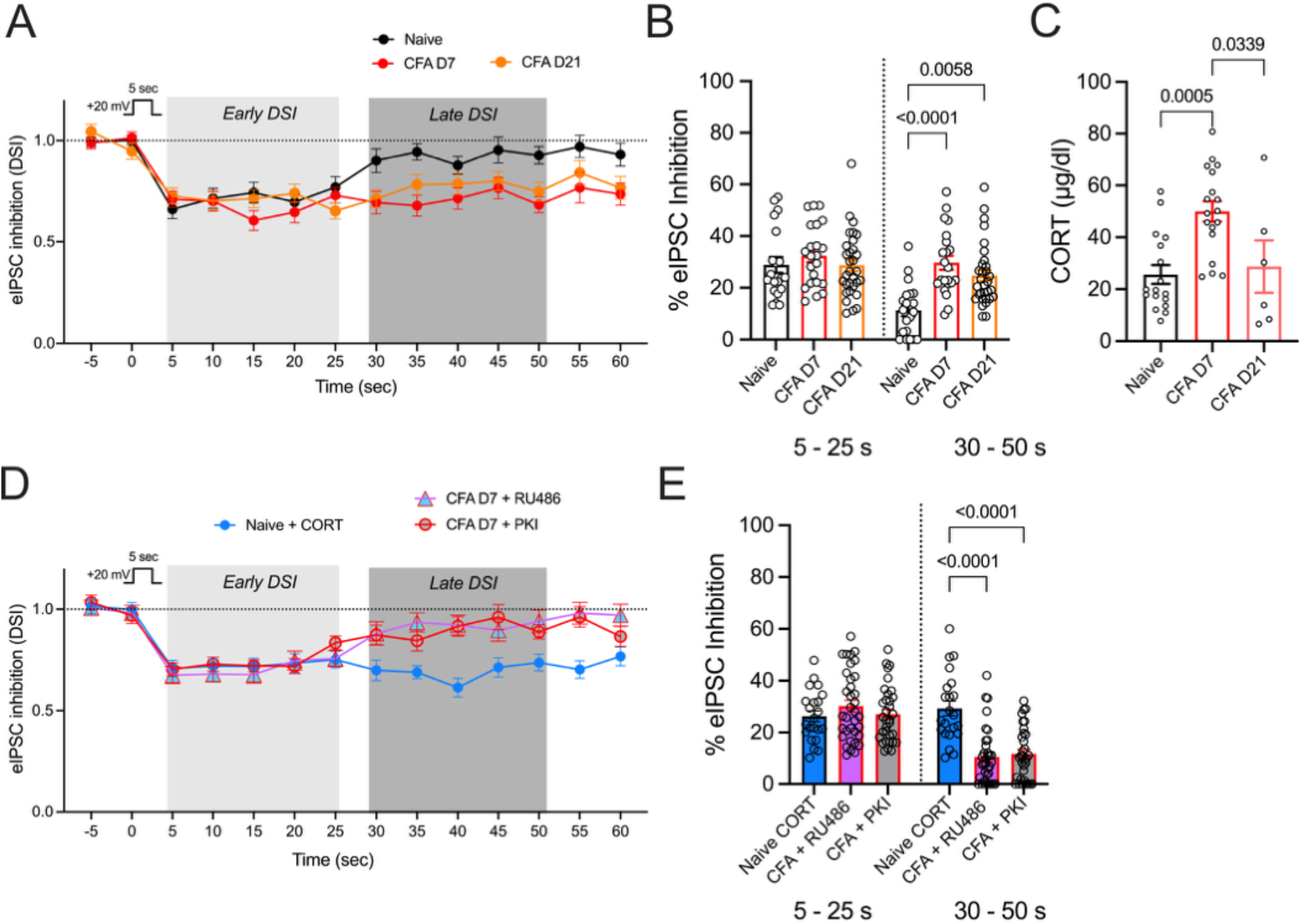
Endocannabinoid function in the vlPAG is enhanced, and regulated by CORT in persistent inflammation. A. The DSI protocol consists of a strong depolarizing step (DSI; +20 mV, 5 s) that induces a transient inhibition of GABA release in recordings from naïve rats that lasts ∼30 s. In CFA rats, the same protocol induces a prolonged DSI at both D7 and D21 post-CFA injections. B. Summary of average % inhibition during the early phase (5 – 25 s) and late phase (30 – 50 s). CFA treatment prolongs DSI up to D21 (All DSI slice treatments are analyzed together (see Table 1) but select comparisons are shown in graphs; Two way RM ANOVA, Time X Treatment, F_8,199_ = 9.59, p < 0.0001, Tukey’s multiple comparisons on graph). C. CORT levels measured in plasma (One way ANOVA, F_2,38_ = 9.40, p = 0.0005, Tukey’s multiple comparisons shown on graph). D. CORT (1 µM) superfused over slices from naïve rats also prolongs DSI. The prolonged phase is blocked by superfusion of RU486 over the slice and PKI in the recording pipette. E. Summary of early and late phase DSI in conditions in panel D. Tukey’s multiple comparisons on graph.

We considered membrane-associated glucocorticoid receptors (GRs) as one possible mechanism for this inflammation-associated increase in endocannabinoid function in the vlPAG. CORT-mediated activation of GRs stimulate synthesis of 2-AG to activate CB1R signaling [25,27,28,52]. Indeed, plasma CORT levels were increased at day 7, and returned to baseline levels by day 21 after CFA injection (Fig. 3C). Exogenous CORT (1 µM) superfusion over slices from naïve rats prolonged DSI, mimicking the prolongation seen in CFA-treated animals, whereas RU486 (5 µM), an inhibitor of GRs, interfered with the prolonged DSI in slices from CFA-treated animals (Fig. 3D,E). Further evidence that GR activation plays a role in the prolonged DSI response came from blocking PKA, since CORT-stimulated 2-AG signaling in the vlPAG is known to be PKA-dependent [27]. Blocking PKA with the peptide inhibitor PKI (300 nM in the recording pipette) readily abolished the late inhibition without affecting early DSI in slices taken at day 7 post-CFA (Fig. 3D,E). Viewed collectively, these observations indicate that increased CORT contributes to potentiated endocannabinoid function in persistent inflammation. It should be noted that RU486 is also a potent inhibitor of progesterone receptors; however, activation of progesterone receptors is known to increase the breakdown of AEA [53], an opposite effect to the one described here.

**Table 1.**

| Two-way RM ANOVA: DSI data |  |  |  |  |  |
| --- | --- | --- | --- | --- | --- |
| ANOVA table | SS | DF | MS | F (DFn, DFd) | P value |
| Time x Treatment | 6698 | 8 | 837.2 | F (8, 199) = 9.593 | P<0.0001 |
| Time | 5325 | 1 | 5325 | F (1, 199) = 61.01 | P<0.0001 |
| Treatment | 18795 | 8 | 2349 | F (8, 199) = 10.69 | P<0.0001 |
| Subject | 43714 | 199 | 219.7 | F (199, 199) = 2.517 | P<0.0001 |
| Residual | 17367 | 199 | 87.27 |  |  |
| Tukey's multiple comparisons test | Summary |  | Adjusted P Value |  |  |
| 5-25 |  |  |  |  |  |
| Naive con vs. Naive CORT | ns |  | 0.9992 |  |  |
| Naive con vs. CFA con | ns |  | 0.9912 |  |  |
| Naive con vs. CFA + CORT | **** |  | <0.0001 |  |  |
| Naive con vs. CFA d21 | ns |  | >0.9999 |  |  |
| Naive con vs. CFA d21 + CORT | ns |  | 0.09 |  |  |
| Naive con vs. CFA + RU486 | ns |  | >0.9999 |  |  |
| Naive con vs. CFA + CORT + Cpd101 | ns |  | 0.9398 |  |  |
| Naive con vs. CFA + PKI | ns |  | 0.9999 |  |  |
| Naive CORT vs. CFA con | ns |  | 0.7858 |  |  |
| Naive CORT vs. CFA + CORT | **** |  | <0.0001 |  |  |
| Naive CORT vs. CFA d21 | ns |  | 0.9989 |  |  |
| Naive CORT vs. CFA d21 + CORT | ns |  | 0.3014 |  |  |
| Naive CORT vs. CFA + RU486 | ns |  | 0.9666 |  |  |
| Naive CORT vs. CFA + CORT + Cpd101 | ns |  | 0.5841 |  |  |
| Naive CORT vs. CFA + PKI | ns |  | >0.9999 |  |  |
| CFA con vs. CFA + CORT | **** |  | <0.0001 |  |  |
| CFA con vs. CFA d21 | ns |  | 0.9768 |  |  |
| CFA con vs. CFA d21 + CORT | ** |  | 0.0056 |  |  |
| CFA con vs. CFA + RU486 | ns |  | 0.9992 |  |  |
| CFA con vs. CFA + CORT + Cpd101 | ns |  | >0.9999 |  |  |
| CFA con vs. CFA + PKI | ns |  | 0.8211 |  |  |
| CFA + CORT vs. CFA d21 | **** |  | <0.0001 |  |  |
| CFA + CORT vs. CFA d21 + CORT | ns |  | 0.312 |  |  |
| CFA + CORT vs. CFA + RU486 | **** |  | <0.0001 |  |  |
| CFA + CORT vs. CFA + CORT + Cpd101 | **** |  | <0.0001 |  |  |
| CFA + CORT vs. CFA + PKI | **** |  | <0.0001 |  |  |
| CFA d21 vs. CFA d21 + CORT | ns |  | 0.054 |  |  |
| CFA d21 vs. CFA + RU486 | ns |  | >0.9999 |  |  |
| CFA d21 vs. CFA + CORT + Cpd101 | ns |  | 0.8823 |  |  |
| CFA d21 vs. CFA + PKI | ns |  | 0.9999 |  |  |
| CFA d21 + CORT vs. CFA + RU486 | * |  | 0.0142 |  |  |
| CFA d21 + CORT vs. CFA + CORT + Cpd101 | ** |  | 0.0026 |  |  |
| CFA d21 + CORT vs. CFA + PKI | ns |  | 0.1437 |  |  |
| CFA + RU486 vs. CFA + CORT + Cpd101 | ns |  | 0.9801 |  |  |
| CFA + RU486 vs. CFA + PKI | ns |  | 0.9822 |  |  |
| CFA + CORT + Cpd101 vs. CFA + PKI | ns |  | 0.6137 |  |  |
| 30-50 |  |  |  |  |  |
| Naive con vs. Naive CORT | *** |  | 0.0001 |  |  |
| Naive con vs. CFA con | **** |  | <0.0001 |  |  |
| Naive con vs. CFA + CORT | ns |  | 0.9783 |  |  |
| Naive con vs. CFA d21 | ** |  | 0.0058 |  |  |
| Naive con vs. CFA d21 + CORT | ns |  | >0.9999 |  |  |
| Naive con vs. CFA + RU486 | ns |  | >0.9999 |  |  |
| Naive con vs. CFA + CORT + Cpd101 | * |  | 0.0112 |  |  |
| Naive con vs. CFA + PKI | ns |  | >0.9999 |  |  |
| Naive CORT vs. CFA con | ns |  | >0.9999 |  |  |
| Naive CORT vs. CFA + CORT | **** |  | <0.0001 |  |  |
| Naive CORT vs. CFA d21 | ns |  | 0.9251 |  |  |
| Naive CORT vs. CFA d21 + CORT | *** |  | 0.0006 |  |  |
| Naive CORT vs. CFA + RU486 | **** |  | <0.0001 |  |  |
| Naive CORT vs. CFA + CORT + Cpd101 | ns |  | 0.9933 |  |  |
| Naive CORT vs. CFA + PKI | **** |  | <0.0001 |  |  |
| CFA con vs. CFA + CORT | **** |  | <0.0001 |  |  |
| CFA con vs. CFA d21 | ns |  | 0.8818 |  |  |
| CFA con vs. CFA d21 + CORT | *** |  | 0.0004 |  |  |
| CFA con vs. CFA + RU486 | **** |  | <0.0001 |  |  |
| CFA con vs. CFA + CORT + Cpd101 | ns |  | 0.986 |  |  |
| CFA con vs. CFA + PKI | **** |  | <0.0001 |  |  |
| CFA + CORT vs. CFA d21 | *** |  | 0.0003 |  |  |
| CFA + CORT vs. CFA d21 + CORT | ns |  | 0.9976 |  |  |
| CFA + CORT vs. CFA + RU486 | ns |  | 0.9881 |  |  |
| CFA + CORT vs. CFA + CORT + Cpd101 | *** |  | 0.0007 |  |  |
| CFA + CORT vs. CFA + PKI | ns |  | 0.9418 |  |  |
| CFA d21 vs. CFA d21 + CORT | * |  | 0.0151 |  |  |
| CFA d21 vs. CFA + RU486 | *** |  | 0.0002 |  |  |
| CFA d21 vs. CFA + CORT + Cpd101 | ns |  | >0.9999 |  |  |
| CFA d21 vs. CFA + PKI | ** |  | 0.0011 |  |  |
| CFA d21 + CORT vs. CFA + RU486 | ns |  | >0.9999 |  |  |
| CFA d21 + CORT vs. CFA + CORT + Cpd101 | * |  | 0.0204 |  |  |
| CFA d21 + CORT vs. CFA + PKI | ns |  | >0.9999 |  |  |
| CFA + RU486 vs. CFA + CORT + Cpd101 | *** |  | 0.0009 |  |  |
| CFA + RU486 vs. CFA + PKI | ns |  | >0.9999 |  |  |
| CFA + CORT + Cpd101 vs. CFA + PKI | ** |  | 0.0039 |  |  |

### GRs contribute to resolution of hyperalgesia

Activation of GRs leads to enhanced CB1R signaling in vlPAG; therefore we predicted that blocking GRs would mimic the behavioral effects of blocking CB1Rs. CFA-treated animals subjected to daily injection of RU486 (5 mg/kg, IP) for up to 10 days exhibited hypersensitivity compared to vehicle-treated animals when tested on day 7 and day 10 (Fig. 4A,B). As in experiments with RIM above, in a separate group of animals, RU486 given only once, at a time point at which hypersensitivity had returned to baseline at D21, reinstated hypersensitivity (Fig. 4C,D). By contrast with these effects in CFA-treated rats, systemic administration of RU486 had no effect on nociception in naïve animals (Supp. Fig. 2). Thus, systemic block of GRs unmasks ongoing hyperalgesia in CFA-treated animals, an effect that parallels hyperalgesia with CBR1 antagonism with RIM injections. After daily injections of RU486 in CFA-treated rats, the late phase of DSI was blocked without any impact on the early phase DSI (Fig. 4E,F). Interestingly, CORT superfusion effectively prolonged DSI similar to the effects observed in naïve rats on day 11 post-CFA in rats treated with daily RU486 injections (Fig. 4G, H). These results indicate that both GRs and CB1Rs were functional in rats following treatment with CFA and RU486 for 10 days.

**Figure 4.**
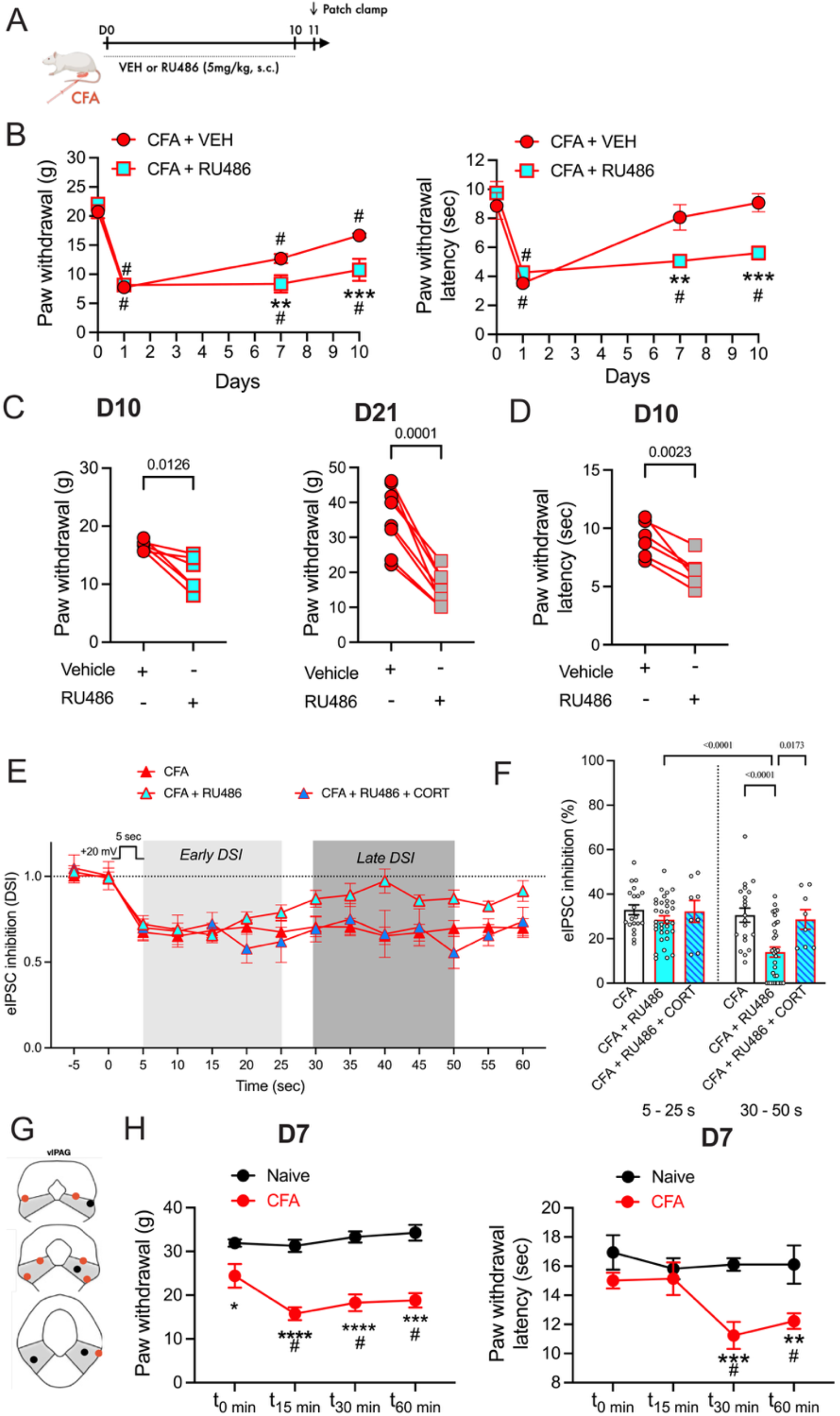
GRs contribute to resolution of hyperalgesia. A. Schematic showing daily injections (SC) of vehicle (VEH) or rimonabant (RIM: 3mg/kg) for 10 days after CFA injections into a hind-paw at D0. B. Daily injections of RU486 (5 mg/kg) in CFA-treated rats reduced recovery from hyperalgesia at D7 and D10 (Two-way RM ANOVA, *Mechanica*l, Time X Treatment, F_3,30_ = 5.505, p = 0.0039, Tukey’s multiple comparisons, CFA compared to VEH, **p = 0.0074, ***p = 0.0004, # Compared to D0 for VEH and CFA; *Thermal*, Time X Treatment, F_3,30_ = 7.781, p = 0.0005, Tukey’s multiple comparisons, CFA compared to VEH, **p = 0.0018, ***p = 0.0004, # compared to D0 for VEH and CFA). C. Rats that received CFA/VEH were given a single injection of RU486 on D10, then tested for mechanical thresholds (paired t-test, t_5_ = 3.801, p = 0.013). A different group of animals were treated with CFA for 21 days and tested with RU486 (Paired t-test, t_7_ = 7.764, p = 0.0001). D. Thermal thresholds were also tested on D10, paired t-test, t_5_ = 5.696, p = 0.0023). E. In the rats treated with CFA and RU486, patch-clamp experiments on D11 were done to test for DSI. The late phase of DSI was reduced in the rats that received daily injections of RU486 and DSI was similar to that observed in slices from naïve rats. Superfusion of exogenous CORT (1 µM) prolonged DSI similar to effects in naïve rats. F. Summary of the early and late phase DSI (Two way RM ANOVA, Time X Treatment, F_2,58_ = 7.903, p = 0.001, Tukey’s multiple comparisons on graph). G,H. Microinjections of RU486 (3 µg/0.4 µL) into the vlPAG also reduce pain thresholds on D7 in CFA-treated rats (Two-way RM ANOVA, *Mechanical*, Time X Treatment, F_3,30_ = 3.274, p = 0.035, Tukey’s multiple comparisons, CFA compared to VEH, *p = 0.013, ****p <0.0001; # compared to D0 for CFA, p < 0.05. *Thermal*, Main Effect of Treatment, F_1,9_ = 10.11, p = 0.011, Tukey’s multiple comparisons, CFA compared to VEH, **p = 0.0056, ***p = 0.0007, # compared to D0 for CFA).

Finally, if these effects of RU486 were mediated by GRs in the vlPAG, we hypothesized that focal microinjection of RU486 in the PAG would potentiate hyperalgesia in CFA-treated rats. Rats were injected and tested at day 7 (Fig. 4G,H). RU486 microinjections (3 µg/0.4 µl) led to a decrease in mechanical and thermal thresholds in CFA-treated, but not naïve, rats over 60 min. Thus, GRs localized to the vlPAG contribute to anti-hyperalgesia and recovery following CFA-induced inflammation.

### Increased endocannabinoid levels prime CB1Rs for desensitization

These data indicate that endocannabinoids can be recruited by glucocorticoids in the vlPAG to limit behavioral hypersensitivity and contribute to recovery from hyperalgesia in animals subject to persistent inflammation. However, dose-response curves for glucocorticoids often form an inverted U, with reduced or no effects at higher doses [54,55]. Moreover, CBR1s are themselves known to desensitize, especially in stress conditions [56]. This raises the possibility that high levels of CORT and associated endocannabinoid action would not have a restorative function, and could even contribute to prolonged hypersensitivity and pain outlasting the period of injury and inflammation.

As shown in previous figures, *endogenous* CORT acting on GRs contributes to enhanced endocannabinoid function after CFA that is reversed by a GR antagonist (Fig. 4E,F above), and addition of *exogenous* CORT (1 µM) to the slice potentiates the DSI response in slices from naïve animals (Fig. 3D, above). By contrast, both the early and late DSI responses were eliminated by exogenous CORT in CFA-treated animals (Fig. 5A). The loss of the CORT response in CFA-treated animals reflected prolonged exposure to endogenous glucocorticoid agonism of GRs, since daily treatment with the GR inhibitor RU486 for 10 days restored the response to exogenous CORT (Fig. 4E,F).

**Figure 5.**
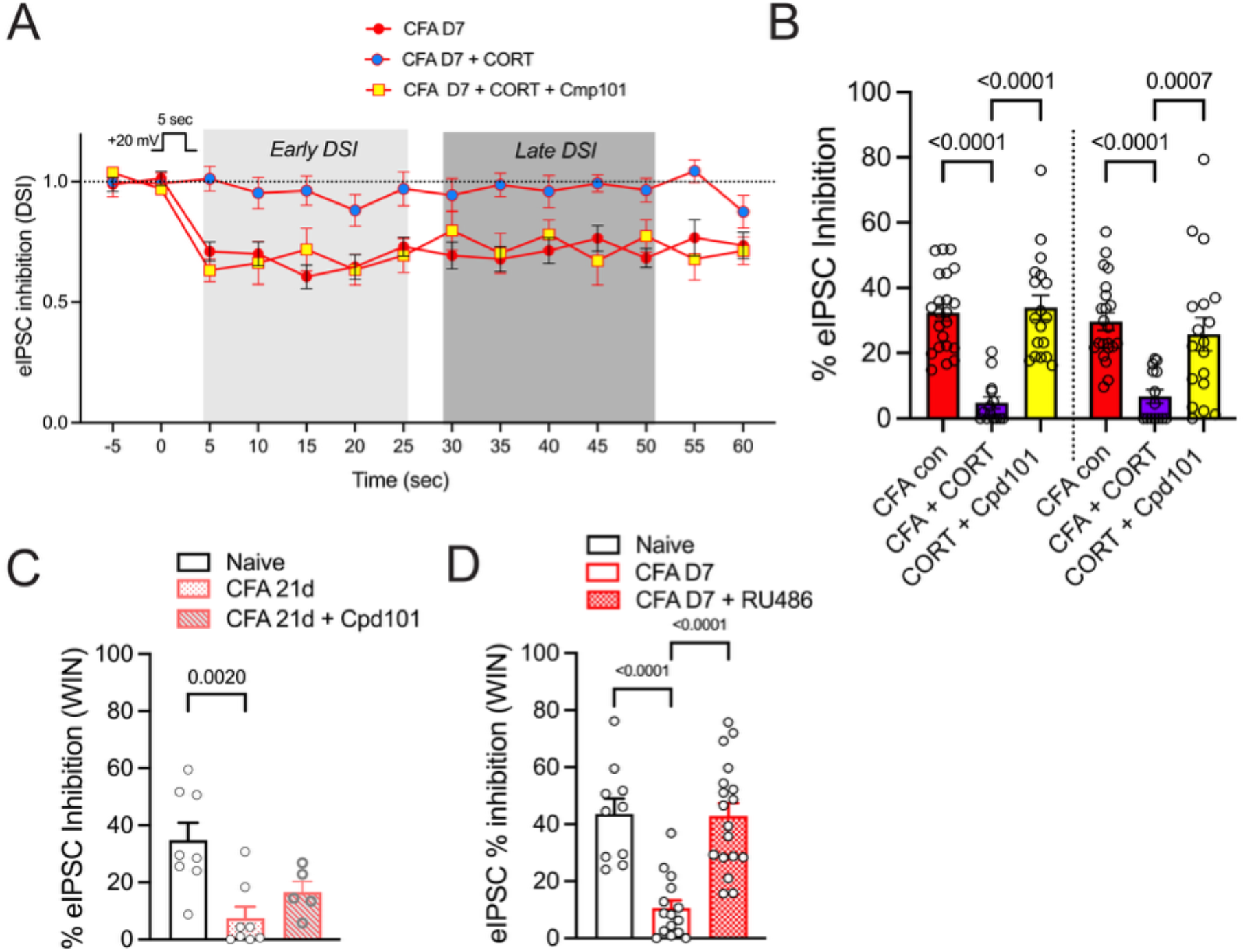
Increased endocannabinoid levels prime CB1Rs for desensitization. A. Superfusion of CORT (1 µM) in naïve rats produces prolonged DSI (Fig. 3D) but both early and late phase DSI is absent in the presence of CORT in CFA-treated rats. Incubation of slices in the GRK2/3 inhibitor Cpd 101 (1 µM) was sufficient to recover DSI in the presence of CORT in recordings from CFA-treated rats indicating that CORT desensitized CB1Rs. B. Summary of DSI in panel A analyzed with all DSI data in Fig. 3 (see Table 1). Tukey’s multiple comparisons shown on graph. C. The effect of WIN55,212-2 (WIN; 3 µM) to inhibit eIPSCs was reduced in CFA rats at D21 compared to naïve rats and partially reversed in the presence of Cpd101 (1 µM). One way ANOVA, F_2,18_ = 8.49, p = 0.003, Tukey’s multiple comparisons on graph. D. The effect of WIN was also reduced at D7 post-CFA and superfusion of RU486 (5 µM) was sufficient to recover the effect of WIN. One way ANOVA, F_2,40_ = 20.34, p < 0.0001, Tukey’s multiple comparisons on graph.

We next considered the possibility that the loss of DSI in vlPAG in the presence of exogenous CORT was due to desensitized CB1Rs. We used Compound 101 (Cpd101, 1 µM), an inhibitor of GRK2/3, a kinase involved in GPCR desensitization[34,57]. Cpd101 recovered both the early and late phase DSI responses in the presence of CORT in CFA-treated rats (Fig. 5A,B). Several additional experiments using the CB1R agonist WIN55,212-2 (WIN) further confirmed that the loss of DSI was explained by desensitized CB1Rs. First, our prior studies showed that WIN inhibition of GABAergic IPSCs was reduced in CFA-treated rats at D7 and Cpd 101 restored WIN inhibition[34]. Here, we confirmed that WIN inhibition was attenuated at day 21 post-CFA with some recovery in the presence of Cpd101 (Fig. 5C). These results indicate that *exogenous* CB1R agonists rapidly desensitize CB1Rs in inflamed rats. However, in rats treated daily with RU486, the effect of WIN was the same as in naïve rats (Fig. 5D), indicating that treatment with RU486 protects CB1Rs from overstimulation.

## Discussion

Here, we provide evidence that the endocannabinoid system is critical for the resolution of hyperalgesia associated with persistent inflammation. Endocannabinoid-dependent latent sensitization is a process mediated by ongoing GR activity in the vlPAG. We also find that, in the context of inflammation, exogenous CORT and CB1R agonists induce desensitization of CB1Rs. Thus, the endocannabinoid system is tightly regulated, creating a narrow therapeutic window for cannabinoid drugs in the context of inflammatory pain.

Changes in endocannabinoid levels during the time course of CFA-induced hyperalgesia were assessed in two ways. First, we measured endocannabinoids in bulk brain tissue using sensitive HPLC mass spectrometry methods. AEA levels increased on day 1 but decreased over the time course of CFA inflammation. In contrast, 2-AG levels were not significantly increased until day 21, with a trend toward increased levels by day 7. Second, we measured endocannabinoid activity at CB1Rs using whole-cell patch-clamp electrophysiology and the DSI protocol which maintains integrity of vlPAG synapses. In these experiments we observed that DSI stimulated CB1R-dependent inhibition of GABA release. DO34, an antagonist of diacylglycerol lipase (DAGL), the enzyme that synthesizes 2-AG, blocks CB1R inhibition of GABA release in the vlPAG in both naïve and CFA-treated rats [27,34] indicating that 2-AG synthesis is required for CB1R-mediated inhibition of GABAergic eIPSCs. Notably, elevation of 2-AG in the vlPAG persists until at least day 21, when hyperalgesia had resolved. This result indicated that endocannabinoids may be a mechanism of latent sensitization in the vlPAG by masking hyperalgesia, similar to the effects of endogenous opioids in the spinal cord [58]. We considered the fact that 2-AG has other known targets, including positive allosteric modulation of GABA_A_ receptors [59,60]. However, this activity would result in an opposite effect, an apparent decrease in CB1R inhibition of GABA release.

To confirm that 2-AG activation of CB1Rs was involved in masking hyperalgesia, we tested the effects of blocking CB1Rs throughout the time course of CFA inflammation. We observed that daily injections of RIM were sufficient to maintain the hyperalgesia through day 10 post-CFA. In fact, a single injection of RIM on day 10 or day 21 in CFA-treated rats produced hyperalgesia and reversed any recovery that had occurred by that day. These data indicate that 2-AG acts at CB1Rs in an ongoing manner up to 21 days post-CFA. This finding also indicates that CB1Rs are functional throughout the time course of CFA-induced inflammation.

We were interested in understanding how CFA injections into the hindpaw and inflammation resulted in long-lasting increases in 2-AG levels in the vlPAG. Based on previous studies showing that CORT activation of GRs can stimulate 2-AG synthesis in many brain regions [52,61], including in the vlPAG [27,62], we hypothesized that CORT could act as a key upstream regulator. Exogenous CORT superfusion over slices from naïve rats produced a prolonged DSI that resembled results from recordings from CFA-treated rats. Moreover, the GR antagonist RU486 blocked the prolonged late phase DSI without affecting the initial phase. Further support that GR signaling is critical for the prolonged phase of 2-AG signaling are the results where the peptide inhibitor of PKA, PKI, applied directly to the recorded neuron through the recording pipette, also blocked the late phase of DSI in recordings from CFA-treated rats. The 2-AG synthesizing enzyme DAGL is activated by PKA phosphorylation [63,64] suggesting that there is a common PKA-dependent mechanism for CORT and inflammation-induced plasticity in DSI experiments [27,34]. Behavioral studies with RU486 administered both systemically and directly into the vlPAG indicate that activation of GRs in the vlPAG play an important role in the resolution of hyperalgesia mediated by CFA. The data also suggest that GRs are critical for endocannabinoid-dependent latent sensitization.

One unexpected finding was that CORT superfusion over slices from CFA-treated rats resulted in a loss of DSI, both the early and late phase DSI. Given that we had established that the early phase of DSI was not dependent on GR activation or PKA activity, but was dependent on 2-AG synthesis, we knew that the early and late phases of DSI were mediated by separate cellular signaling pathways that converged on 2-AG signaling to CB1Rs. We tested the possibility that CB1Rs desensitized in the presence of inflammation plus exogenous CORT and found that the inhibitor of GRK2/3-mediated desensitization was sufficient to restore DSI. This result was consistent with our prior findings that blocking the breakdown of 2-AG in slices from CFA-treated rats also resulted in desensitization of CB1Rs[34]. Interestingly, plasma levels of CORT have returned to normal by day 21, so either tissue levels of CORT are still elevated and sufficient to activate GRs or GRs are activated by metabolites of CORT that are agonists of GRs [65].

Our findings highlight a paradox: while CORT-induced 2-AG signaling may initially be protective, persistent elevation of the endocannabinoid system risks CB1R desensitization, ultimately reducing the analgesic efficacy of exogenous direct or indirect CB1R agonists. This is an interesting observation given that presynaptic GPCRs do not readily desensitize [34,66,67]. Presynaptic CB1Rs in the vlPAG are resistant to exogenous cannabinoid agonists for more than 5 hours [34] and even after 21 days of elevated 2-AG levels, as shown in this paper. Chronic or repeated cannabinoid agonists, however, have documented significant CB1R desensitization, receptor internalization, uncoupling from Gi/o proteins, and reduced signaling efficacy across multiple brain regions[68-70]. For example, CB1R internalization has been identified as a central mechanism of agonist-induced desensitization[71], while genetic deletion of MAGL increases 2-AG levels and leads to brain region-specific desensitization of CB1R Gi/o signaling and reduced agonist responses [72,73]. In our persistent inflammation model, preventing CB1R desensitization (GRK2/3 inhibition with Cpd101) restores CB1R-mediated suppression of GABA release in vlPAG, directly linking receptor responsiveness to preserved short-term plasticity and descending pain modulation.

From a translational perspective, a key challenge of cannabinoid-based analgesics is the development of tolerance, similar to opioid drugs [69,74-76], thereby limiting their long-term efficacy in pain management. Administration of WIN confirmed the loss of CB1R-mediated anti-hyperalgesia in CFA-treated animals, consistent with CB1R desensitization. RU486 blockade of GR signaling during CFA inflammation preserves analgesic effects of WIN in both thermal and mechanical modalities. Thus, GR antagonism prevents CB1R overstimulation and desensitization, maintaining cannabinoid analgesic efficacy. However, GR antagonism alone sustains pain, highlighting the narrow therapeutic window for CB1R agonists in the context of inflammation and the need for balanced strategies. Furthermore, endocannabinoids are both increased and decreased in the presence of inflammation and neuropathic pain in humans and animal models depending on the tissue measured and pain condition[44,77-80]. It is clear that it is not sufficient to measure only levels of endocannabinoids but also assessing whether the receptors are functional is critical to develop effective new pain therapeutics targeting the cannabinoid system.

In conclusion, CORT provides a protective mechanism during persistent inflammation but the context of inflammation also renders CB1Rs more vulnerable to desensitization or “primed” for desensitization by exogenous and endogenous cannabinoid agonists. Therapeutically, combining CB1R agonists with GR modulation, or developing allosteric modulators of CB1R that do not promote CB1R desensitization, may broaden the therapeutic window for cannabinoid-based drugs to preserve analgesia while limiting tolerance.

## Author contributions

S.L.I., B.C. and C.A.B. conceived the research and designed electrophysiological experiments, B.C., C.A.B., L.C.P., C.M.D., and D.C.J. performed and analyzed electrophysiology and behavioral experiments, Je.K and J.K. designed LCMS experiments and I.G. performed LCMS experiments, B.C., M.M.H., and S.L.I. wrote and editedthe manuscript.

## Data Availability Statement

Data is available upon request. Please contact Susan Ingram, PhD, Department of Anesthesiology, University of Colorado Anschutz, 12700 E 19th Ave, P15-7480D, Aurora CO 80045.

## Supplementary information

## Supplemental Detailed Materials and Methods

**Supp. Table 1**. Statistical analysis of DSI data.

### Drugs

WIN55,212-2 (Caymen Chemicals Ann Arbor MI, USA), SR141716A (RIM; Cayman Chemical), Corticosterone (CORT; Tocris, Minneapolis MN, USA), 11b-(4-dimethyl-amino)-phenyl-17bhydroxyl-17-(1-propynyl)-estra-4,9-dien-3-one (RU486; Tocris) were dissolved in DMSO, aliquoted, and stored at - 20°C. NBQX (Tocris) was dissolved in milliQ water, and stored at 4°C. Compound101 (Cpd101, HelloBio, Bristol UK) was first dissolved in a small amount of DMSO (10% of final volume), sonicated, then brought to its final volume with 20% 2-hydroxypropyl)-b-cyclodextrin (HPCD) and sonicated again to create a 10 mM solution. PKA inhibitor (PKI, Tocris) was used directly in the internal solution in recording electrodes at 0.2 µM.

### vlPAG slice preparation

Slices containing the vlPAG were prepared as previously described [34,40]. Rats were deeply anesthetized with isoflurane (McKesson, Irving TX, USA), and the brain was rapidly removed and placed in ice-cold aCSF cutting buffer containing the following (in mM): 126 NaCl, 21.4 NaHCO3, 22 dextrose, 2.5 KCl, 2.4 CaCl2, 1.2 MgCl2, and 1.2 NaH2PO4 (300 mOsm). Slices containing the vlPAG were cut to a thickness of 220 µm on a vibratome (Leica Microsystems, Deerfield IL, USA) and were transferred to a holding chamber maintained at 32°C. Slices were oxygenated with 95% O_2_/5% CO_2_ until transfer to the recording chamber on an upright microscope (model BX51WI, Evident Scientific, Waltham MA, USA) and superfused with oxygenated aCSF maintained at 32°C.

### Whole-cell patch-clamp recordings

Voltage-clamp recordings (holding potential, -70 mV) were made in whole-cell configuration using an amplifier (MultiClamp 700B, Molecular Devices), sampled at 2 kHz, and digitized at 5 kHz with the Axon Digidata 1550B (Molecular Devices, USA) using Clampex 11.0.3 software (Molecular Devices, USA). Patch-clamp electrodes were pulled from borosilicate glass (diameter, 1.5 mm; WPI) on a two-stage puller (Narishige). Pipettes had a resistance between 2.5 -4 MΩ and were filled with an intracellular pipette solution containing the following (in mM): 140 CsCl, 10 HEPES, 4 MgATP, 3 NaGTP, 1 EGTA, 1 MgCl2, and 0.3 CaCl2 (pH 7.3, 290–300 mOsm). QX314 (100 µM) was added to the internal solution for evoked IPSC (eIPSC) experiments to reduce action potentials in the recording cell. Access resistance was continuously monitored. Recordings in which access resistance changed by 20% during the experiment were excluded from data analysis. A bipolar stimulating electrode (FHC, Bowdoin ME, USA), placed into the vlPAG approximately 200 µm from the recording electrode, was used to deliver 2 ms pulses of 100 µA to 10 mA to evoke IPSCs. A junction potential of 5 mV was corrected during recording. GABAergic eIPSCs were isolated in the presence of glutamate receptor antagonist (NBQX; 5 µM). In experiments using exogenous cannabinoid agonists/antagonists, or GR agonists/antagonists, only one neuron was recorded per slice. After each experiment, the lines were washed with 70% ethanol and then rinsed with milliQ water.

### Endocannabinoid analysis

Quantification of 2-AG and AEA level were performed in the vlPAG, the RVM and the injured paw tissue of naïve and CFA-treated rats. Tissues were removed, quickly frozen in liquid nitrogen and stored at -80ºC degrees until analysis. Approximatelty 50 mg of tissue powder were placed into 1000 µL of acetonitrile/methanol (1/1, v/v) containing an isotope labeled internal standard mix on a scale to record the actual tissue weight. Samples were shaken and centrifuged at 25000xg, 4ºC for 15 minutes. Supernatants were transferred to HPLC sample vials and analyzed using HPLC mass spectrometry as previously described [42-44]. Following shaking and centrifugation, the supernantant was analyzed as described above and protein pellet was dried and reconsituted in detergent buffer to determine the protein content.

### Measurement of plasma CORT levels

Plasma CORT levels were determined using trunk blood immediately after anesthesia at the time of slicing for electrophysiological recordings in the morning to avoid variation due to circadian rhythm. Blood fractionation was achieved by spinning the blood samples at 14000 rpm for 20 min using a centrifuge (Model 5418R, Eppendorf). Supernatant containing plasma was rapidly collected and stored at −80°C until subsequent analysis. All samples were analyzed using the commercially available enzyme-linked immunosorbent assay (ELISA) CORT kit (#EIACORT, ThermoFisher Scientific) according to the manufacturer’s specifications.

### Behavioral studies

**Mechanical nociception** was assessed using an electronic von Frey (Ugo Basile, Gemonio, Italy). Prior to the experiment, rats were acclimatized to a Plexiglas chamber for 15 min (grid: 0.5 cm × 0.5 cm; box: 10 cm × 10 cm × 15 cm) on a raised-mesh metal platform. Responses from each hind paw were measured three times each, and the mean for each paw was calculated as the paw withdrawal threshold (PWT). Baseline responses were measured before CFA injection (D0) and at various time points. All measurements were performed by the same person, blinded to treatment.

**Thermal nociception** was assessed using the Hargreaves apparatus (Plantar Test, Ugo Basile, Gemonio, Italy), which measures paw withdrawal latency (PWL) to radiant heat stimuli. Rats were acclimatized in singular Plexiglas chambers (10 cm × 10 cm × 15 cm) on a glass platform for 15 min. The radiant heat source was applied to the center of the plantar surface of each hind paw with 2 min intervals between each application, and PWL determined as time to retraction or licking of the hind paw. To avoid tissue damage, a cut-off of 25 seconds was used. All trials were performed three times for each hind paw, and the average for each hind paw was calculated as the PWL. All measurements were performed by the same person who was blinded to the treatment.

### vlPAG microinjections

To assess the central effect of RU486 on behavior, rats performed mechanical and thermal nociception tests as previously described, then the animal were anesthetized with vetflurane (3%, Piramal, PA USA) using a R500 Small Animal Anesthesia Machine (RWD, Sugar Land, TX USA). Unilateral microinjection of RU486 (3 µg/kg dissolved in 10% DMSO in saline, 0.4 µl in one minute) into the vlPAG was performed using a stereotaxic apparatus (NeuroSTAR, Germany). The injection cannula was left in place for 2 min to minimize backflow. Immediately after microinjection, wounds were closed and animals were allowed to recover for 5 min. Animals were subjected to the behavioral testing protocols 15, 30 and 60 min post injection.

### Statistical analysis

In all electrophysiological experiments, each dataset included recordings from at least two male and two female rats. For DSI experiments, two separate time windows were analyzed: the average of the first five eIPSCs (0–25 s) following depolarization, and the average of the subsequent five eIPSCs (30–50 s). All analyses were conducted in Graphpad Prism (Graphpad Software). Values are presented as the mean ± SEM, and all data points are shown in bar graphs to illustrate variability. Statistical comparisons were made using t-test or ANOVA, as appropriate. In all summary bar graphs for electrophysiology experiments, each dot represents an individual cell while the numbers in the bars represent the animal number. When post hoc analysis was appropriate, multiple-comparisons tests were performed and specified in the figure legends. Statistical significance was defined as p < 0.05.

## Supplemental figures

**Supplementary Fig 1.**
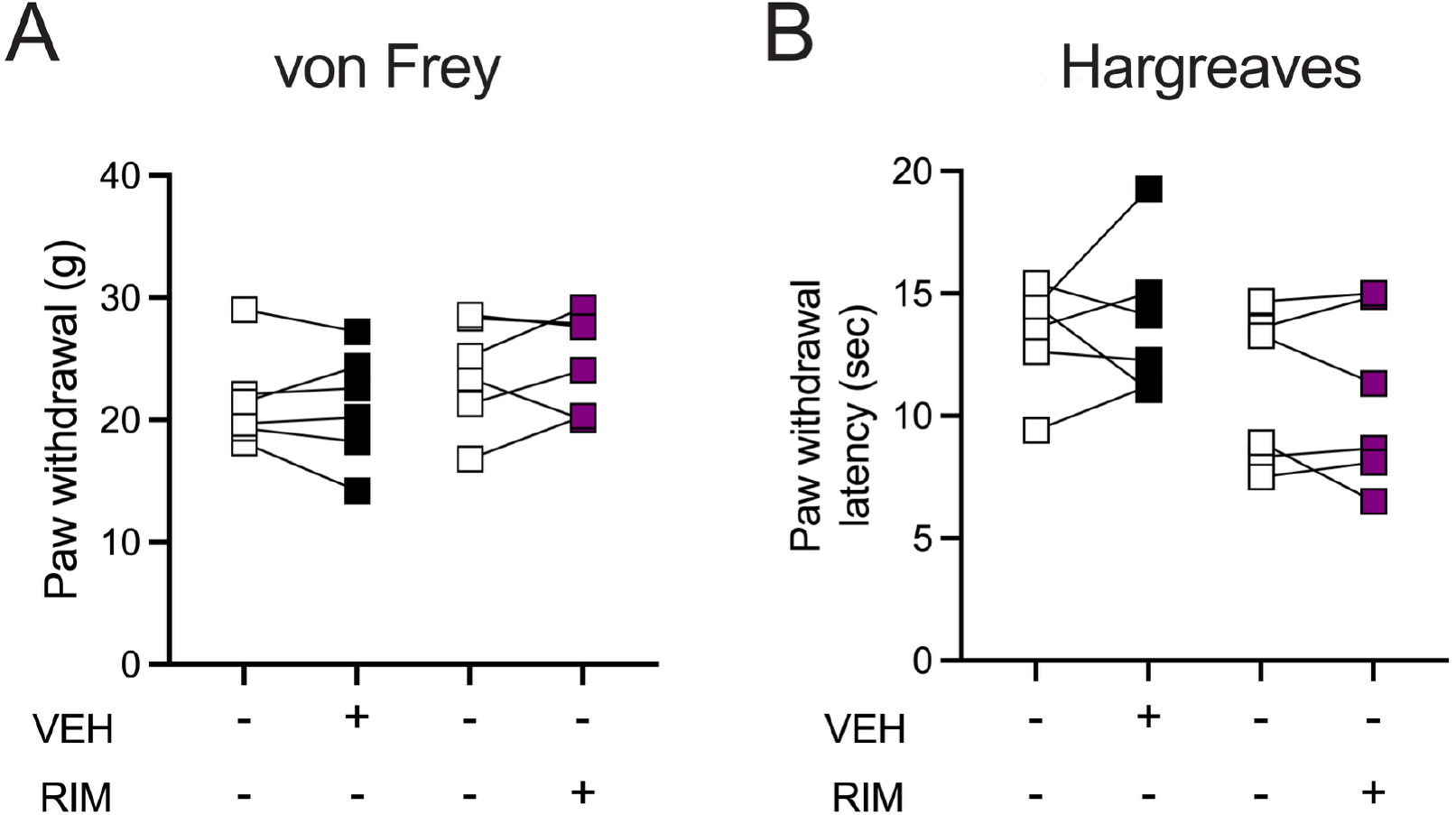
Repeated injections of vehicle or RIM do not affect pain thresholds in naïve rats. A. Naïve rats received daily injections of vehicle or RIM (3 mg/kg, SC) over 10 days and were then tested for mechanical thresholds (Two way ANOVA *ns* for Interaction, Drug or Time). B. Rats were also tested for thermal thresholds (Two way ANOVA *ns* for Interaction, Drug or Time).

**Supplementary Fig 2.**
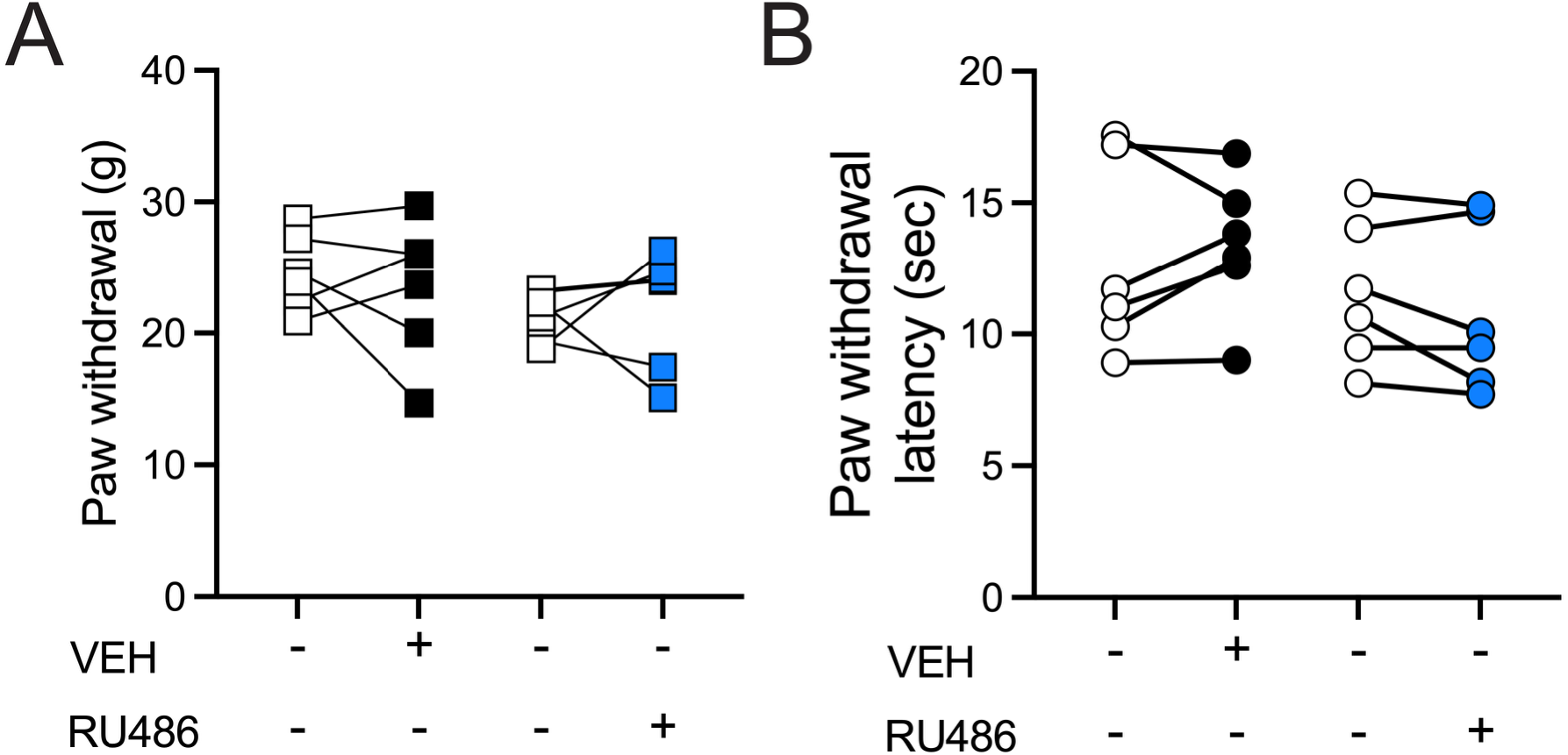
Repeated injections of vehicle or RU486 do not affect pain thresholds in naïve rats. A. Naïve rats received daily injections of vehicle or RU486 (5 mg/kg, SC) over 10 days and were then tested for mechanical thresholds (Two way ANOVA *ns* for Interaction, Drug or Time). B. Rats were also tested for thermal thresholds thresholds (Two way ANOVA *ns* for Interaction, Drug or Time).

